# Statistical characterization of tactile scenes in three-dimensional environments reveals filter properties of somatosensory cortical neurons

**DOI:** 10.1101/2022.08.03.502632

**Authors:** Nadina O. Zweifel, Sara A. Solla, Mitra J. Z. Hartmann

## Abstract

Natural scenes statistics have been studied extensively using collections of natural images and sound recordings. These studies have yielded important insights about how the brain might exploit regularities and redundancies in visual and auditory stimuli. In contrast, natural scenes for somatosensation have remained largely unexplored. Here we use three-dimensional scans of natural and human-made objects to quantify natural scene statistics at the scale of the human fingertip. Using measurements of distance, slope, and curvature from the object surfaces, we show that the first order statistics follow similar trends as have been observed for images of natural and human-made environments. In addition, independent component analysis of curvature measurements reveals Gabor-like basis vectors similar to those found in natural images. A simple neural model using these filters showed responses that accurately capture the statistics of responses in primate primary somatosensory cortex.

## 1. Introduction

An animal’s only access to the environment is through the signals provided by the activation of its sensors as they interact with the external world. These sensory signals typically represent compressed and nonlinear transformations of physical quantities in the environment. Nonetheless, nervous systems can efficiently and reliably infer relevant information about the original environmental variables from the sensory input signals they evoke. It has been hypothesized that these successful inferences are achieved by exploiting regularities and redundancies present both in the environment and in the resulting sensory input signals [1, 2].

Over the last five decades, a large body of research on the statistics of visual and auditory natural scenes has generated strong evidence in support of this hypothesis [3–21]. Statistical models that predict the coding schemes and representations characteristic of cortical neurons (basis functions, basis vectors, receptive fields) have received considerable attention, and different algorithms to fit the models to sensory data have been proposed [4, 5, 14, 22–27]. The underlying assumption of these models is that sensory systems process incoming information by simultaneously activating a small subset of neurons while the majority remain silent. Some modeling approaches directly impose this constraint by maximizing the sparseness of the optimized representations (sparse coding) [22], while others indirectly achieve similar results by maximizing the independence of the predicted basis functions (independent component analysis) [4, 28].

The data used to train these models typically originate from a collection of natural scenes in the form of visual images or sound recordings; these represent the ensemble of inputs that the sensory system may encounter. For images, a large number of small regions within these images is then used to mimic receptor-like sampling of the scene. The basis functions that emerge from the fitted model thus represent the features of the sensory signal encoded by individual cortical neurons; these have been found to be very similar to cortical receptive fields recorded experimentally [4, 22, 28]. The features thus selected can give insight into the neural tuning that gives rise to an efficient code. In vision, these features resemble edge detectors and Gabor filters [22, 26, 28].

In contrast to vision and audition, the natural scene statistics of somatosensation have remained largely unexplored [19–21, 29]. Tactile sensors are unique in that the signals they acquire result from direct physical contact with the environment: it is impossible to “feel” an object without touching it. The interactive nature of touch makes tactile environmental statistics extremely challenging to quantify, as evidenced by the small number of studies that have attempted to measure these parameters *in vivo* [21] or *in silico* [19, 20, 30]. To date, psychophysical, neurophysiological, and self-report experiments have primarily focused on the physical and qualitative dimensions of tactile perception in the environment (e.g., roughness, smoothness, compliance, curvature, etc.) [31–38]. Despite substantial groundwork from these studies, the environmental statistics of somatosensation are still poorly documented [29].

The present study aims to shed light on the environmental statistics of the natural *tactile* scene by quantifying geometrical properties of the natural environment in three-dimensional (3D) space. We focus specifically on three different Cartesian-based metrics: local distance, slope, and curvature. We begin by quantifying the statistics of each of these metrics at the level of individual objects as well as the entire population of objects, and then use independent component analysis (ICA) in two different ways. First, ICA is used to find the leading statistically independent dimensions in local distance measurements, revealing that 95% of the variance in local distance data is predominantly accounted for by curvature. Second, ICA is used to estimate the filter characteristics of somatosensory cortical neurons. This analysis shows that the ICA basis vectors estimated from curvature patches have Gabor-like filter properties and are consistent with orientation sensitivity found in neurons in areas 1 and 3b of primary somatosensory cortex (S1).

## 2. Results

We collected 3D representations of 96 natural objects such as rocks, leaves, bark, fruits, vegetables, etc., as well as 41 human-made objects including scissors, knifes, purses, mugs, etc. (Fig. 1ab). To quantify the local shape of these data, we randomly sampled 100-150 circular surface patches with a radius of 6 mm, similar to the size of a human fingertip [39] (Fig. 1c, *left*). This procedure resulted in a total of 12,309 and 5,788 patches for natural and human-made objects, respectively. For each surface patch we computed maps of distance, slope, and curvature; details for calculating these metrics are found in *Methods.* Briefly, the distance of the surface points was computed with respect to a reference plane (Fig. 1c, *middle*), yielding a circular distance map ***P***(*D*) with a total of 448 points for each patch. From ***P***(*D*) we computed a 421-dimensional slope map ***P***(*S*) using first-order forward difference, and a 384-dimensional curvature map ***P***(*C*) using second-order central difference. Examples of ***P***(*D*), ***P***(*S*), ***P***(*C*) for an example patch are shown on the right in Fig. 1c.

**Fig. 1.**
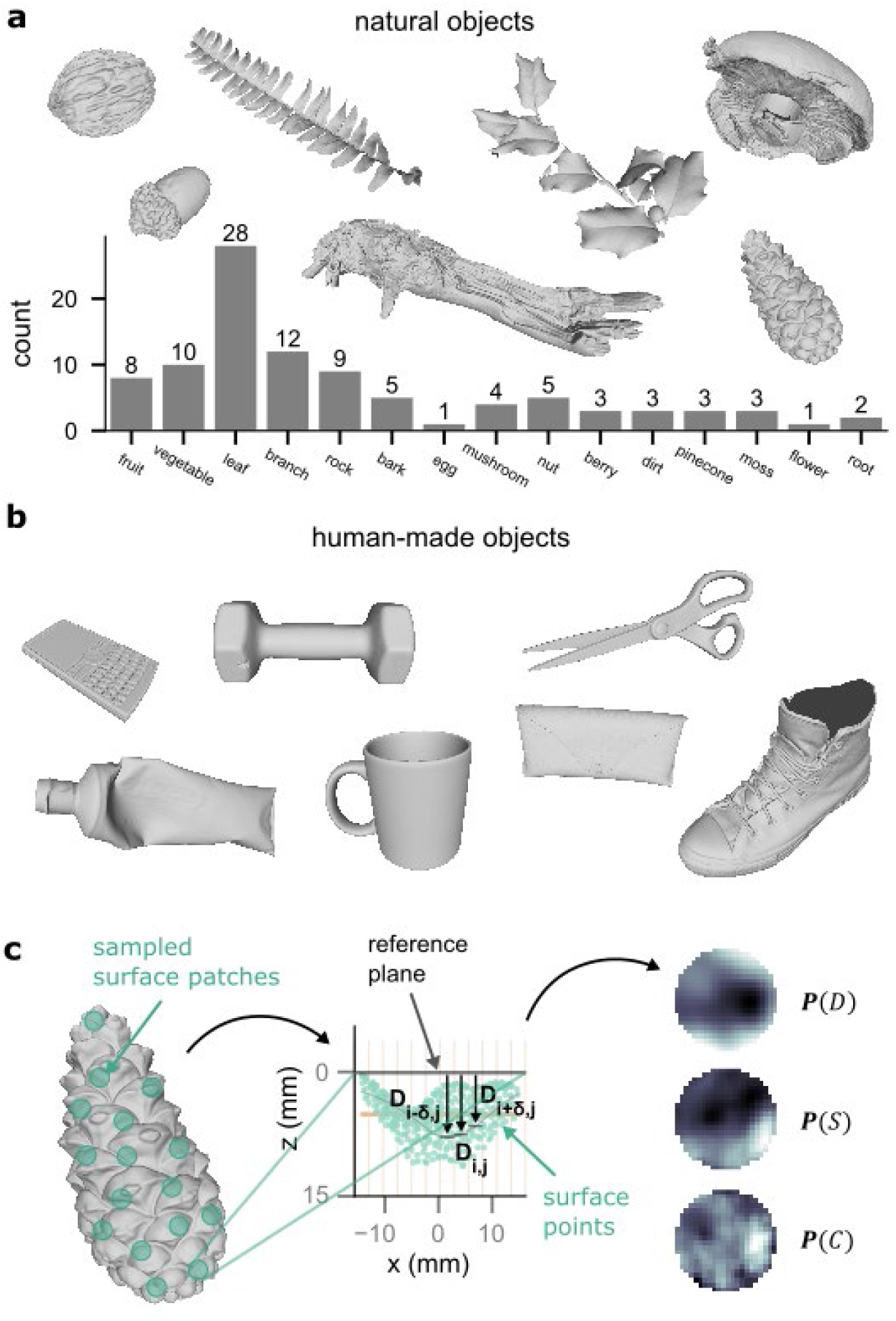
The 3D datasets include a total of 96 natural objects and 41 human-made objects, cleaned and processed following procedures described in *Methods.* **(a)** Examples of 3D scanned natural objects, and a histogram of the number of objects in each of 15 categories. **(b)** Examples of 3D scanned human-made objects. **(c)** To sample the 3D data, approximately 100-150 circular surface patches were randomly selected from each object (left). For each surface patch, the distance of the surface points was measured with respect to a reference plane (middle). From these measurements, a circular distance map ***P***(*D*) of 448 points was created (right). Using forward and central differencing, a 421-dimensional slope map and a 384-dimensional curvature map, ***P***(*S*) and ***P***(*C*), were computed for each patch (see *Methods).*

### 2.1. First-order statistics of natural and human-made objects

We examined the first-order statistics of natural and human-made objects in terms of the local metrics: distance, slope, and curvature (*D, S,* and *C*). Fig. 2a compares the probability distributions of these metrics for two natural objects (bark, leaf) and two human-made objects (mug, purse). The distributions were computed from all patches for each object. As expected, human-made objects tend to have larger flat and low-curvature regions (e.g., the body of the mug); the probability distributions for all three metrics are correspondingly skewed towards zero. At the same time, small distinct regions of human-made objects exhibit higher curvature values (e.g., rim and handle of the mug) that affect the shape of the corresponding distributions. The effect of the rim and handle of the mug is particularly obvious in the distribution of *C,* in which additional modes appear at large positive and negative values. In contrast, the distributions for natural objects are clearly unimodal with a smooth decay towards the tails, indicating that the measured local metrics are more continuous across the surface than for human-made objects.

**Fig. 2.**
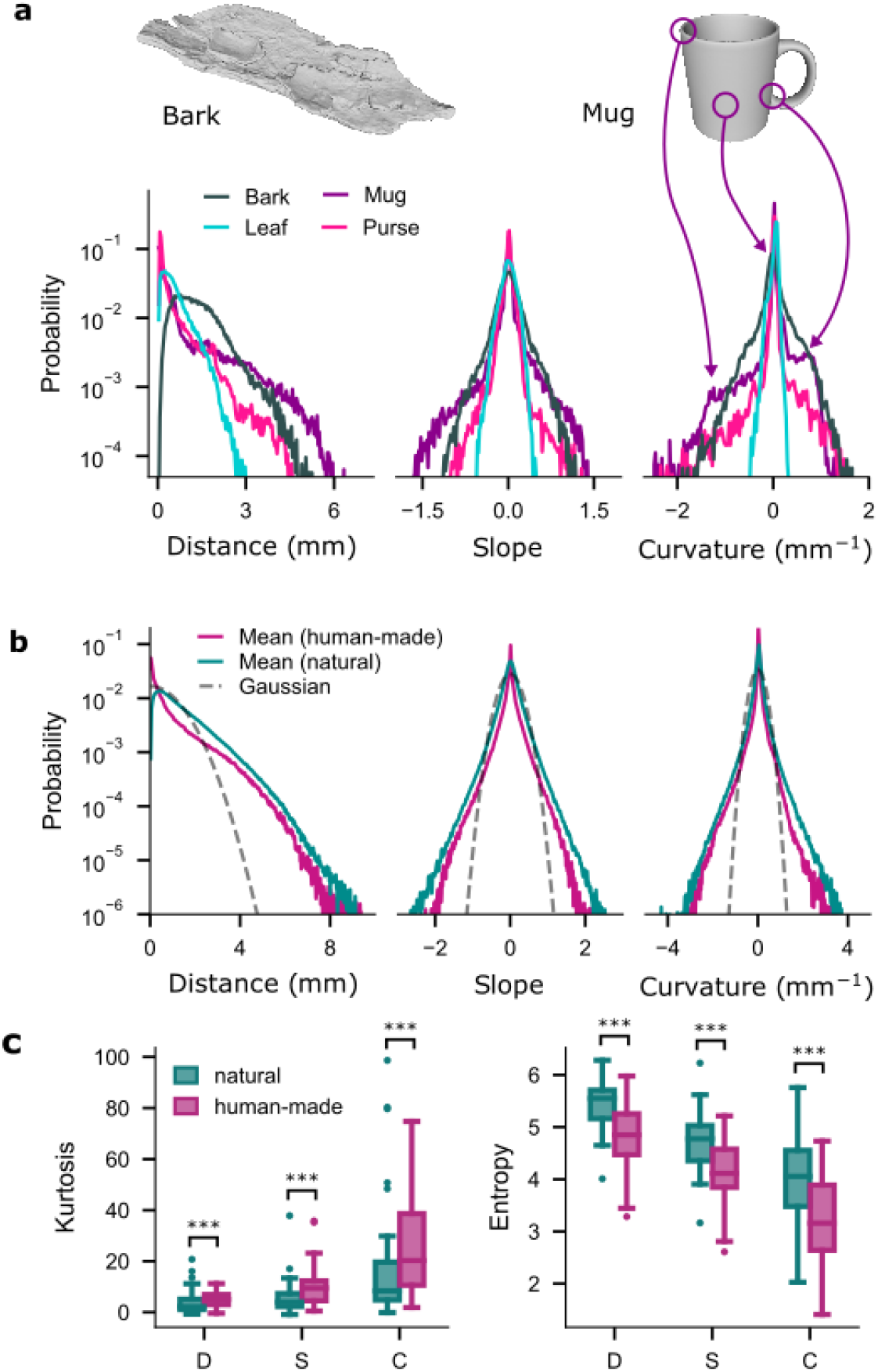
Probability distributions of the three metrics *D, S,* and *C* for both natural and human-made objects. **(a)** Four example objects (two illustrated, not to scale) and their corresponding distributions of *D* (left), S (middle), and *C* (right). Distributions were computed from the ensemble of patches with *R* = 6 mm associated with each object. Note that the vertical axis is logarithmic. **(b)** Average probability distribution of *D* (left), S (middle), and *C* (right) across all sampled patches with *R* = 6 mm for natural (turquoise) and human-made (magenta) objects. The black dashed lines show a zero-mean Gaussian distribution of the same variance for comparison. Only the positive half of the Gaussian is used for distance *D.* **(c)** Distributions of kurtosis (left) and entropy (right, measured in bits) for the distribution for each object class and each metric.

To generalize this analysis, we computed the average probability distributions across all ***P***(·) for each metric (Fig. 2b). Both natural and human-made objects result in super-Gaussian distributions, i.e., they have higher peaks at zero and heavier tails than a Gaussian distribution with the same mean and variance. Although the average distributions for human-made objects smooth over irregularities present in the distributions for individual human-made objects, the average distributions for human-made objects remain heavily skewed towards zero and are significantly different from the distributions for natural objects distance *D* (Kolmogorov-Smirnov test: D = 0.303, p < 0.001), slope S (Kolmogorov-Smirnov test: D = 0.100, p < 0.001), and curvature *C* (Kolmogorov-Smirnov test: D = 0.149, p < 0.001).

Differences between natural and human-made objects can also be quantified in terms of kurtosis: human-made objects result in significantly higher kurtosis values than natural objects (Fig 2c, left) in terms of *D* (Mann-Whitney rank test: U = 1029, p < 0.001), S (Mann-Whitney rank test: U = 892, p < 0.001), and *C* (Mann-Whitney rank test: U = 984, p < 0.001). The average distributions of *D, S,* and *C* have kurtosis values of 2.8, 6.1, and 17.9 for natural objects, and 4.4, 8.9, and 24.3 for human-made objects, respectively. Kurtosis is related to regularity (correlations) in the data and is inversely related to entropy, i.e., high kurtosis values correspond to low entropy, shown in the right panel of Fig. 2c. Entropy is significantly lower for human-made objects than for natural objects for all three metrics *D* (Mann-Whitney rank test: U = 508, p < 0.001), S (Mann-Whitney rank test:, U = 791: p < 0.001), *C* (Mann-Whitney rank test: U = 696, p < 0.001). From an information-theoretical perspective, this result suggests that fewer bits are necessary to encode information about human-made objects than for natural objects when the objects are characterized through the chosen metrics *D, S,* and *C.* In addition, the right plot in Fig. 2c also indicates that entropy decreases with the order of the derivative: *D* results in higher entropy and *C* in lower entropy across individual objects. This result implies that a metric such as *C* may provide more regularities that could potentially be exploited by a sensory system to efficiently encode information about tactile images present in the world.

### 2.2. ICA analysis for the various metrics

The first-order statistics of three local metrics that characterize the shape of *R* = 6mm surface patches: distance, slope, and curvature are highly regular and super-Gaussian, congruent with the statistics observed in natural images [40, 41]. However, it remains unclear which of these metrics may be the best candidate to provide information about the world to a tactile sensory system. To investigate this question, we used an unsupervised learning algorithm, Independent Component Analysis (ICA), to identify the fundamental features of natural data in terms of distance *D*, the least processed metric that characterizes local shape.

As described in [42], ICA is a linear model that maps a data vector *x* consisting of a set of features [*x*_1_ …, *x_n_*] to a set of components *s* = |*s*_1_, …,*s_n_*] through a square mixing matrix *A,* such that the sensory signal *x* can be linearly reconstructed from the components by

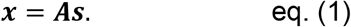

Both the mixing matrix ***A*** and the components s are unknown. The matrix ***A*** is estimated from the statistics of the observed data vectors ***x*** under the assumptions that ***A*** is square, the components s are statistically independent, and ***x*** can be reconstructed linearly [43]. The independent components s are then obtained by using the matrix ***W***, the inverse of ***A***:

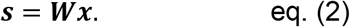

The columns in ***A*** are called basis vectors; each basis vector is associated with one of the components in ***s***. The rows in ***W*** play the roles of filters that act on the data vector ***x*** to obtain the corresponding component [28]. The matrix ***W*** contains the coefficients or weights for each of the features in ***x*** to recover each independent component in ***s***. The set of weights associated with a particular component *s_i_*, which correspond to the 7-th row of ***W***, can thus be interpreted as the receptive field for that component [28]. In contrast, the column basis vectors in ***A*** represent fundamental features that can be linearly combined to compose the observed signal ***x***. The amplitude of each such feature is determined by the coefficients of the corresponding component of ***s***; these components are typically sparsely distributed [28]. There are many different ways to implement ICA; we used one of the most popular and computationally efficient algorithms, FastICA, proposed by Hyvarinen and Oja [43].

In our scenario, ***x*** represents a 448-dimensional vector of distances from a map ***P***(*D*) sampled from a natural object. Examples of ***P***(*D*) used for fitting the ICA model are shown in Fig. 3a. The FastICA algorithm does not sort the estimated basis vectors (columns of A), so we used a sorting algorithm proposed by Cheung and Xu [44] to order the basis vectors according to their contribution to reconstruction accuracy. The first 100 of the sorted basis vectors are shown in Fig. 3b. The cumulative reconstruction accuracy as sorted basis vectors are added is shown in Fig. 3c; this plot indicates that the leading 113 basis vectors suffice to achieve a reconstruction accuracy of 0.95. Based on this result, we determined the filtering properties of the leading 113 basis vectors by convolving each of them with a test image (Fig. 3d, *left inset).* The convolution output allowed for a categorization of the basis vectors into 1^st^, 2^nd^, and 3^rd^-order derivative filters. Examples of each category with the corresponding convolved images are shown in Fig. 3d. Note that 1^st^-order filters, associated with slope, are notably scarce, rising slowly in number as basis vectors are added. Within the first 113 basis vectors, only four 1^st^-order filters were present (Fig. 3e). In contrast, 2^nd^-order filters, associated with curvature, were the most common.

**Fig. 3.**
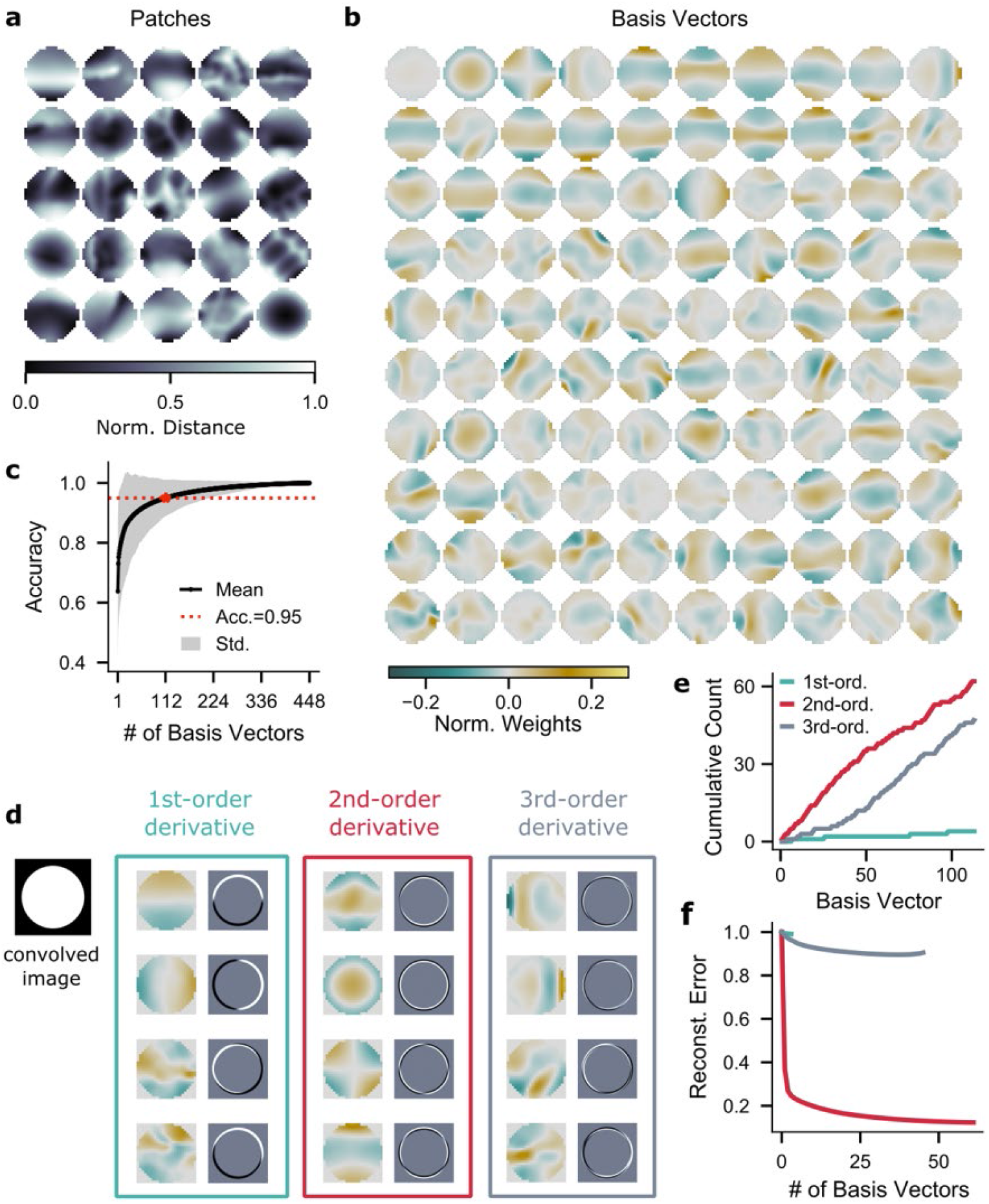
Basis vectors estimated by ICA for surface patches characterized by distance *D* to the reference plane. **(a)** Example surface patches (*R* = 6mm) randomly sampled from natural 3D data that show spatial maps of the distance *D.* Grey scale is normalized to maximum distance within each patch. **(b)** The leading 100 basis vectors (columns of the mixing matrix 4) as estimated by ICA from the distance *D* data. The mean was subtracted from each basis vector, and the norm of each basis vector was scaled to 1. The basis vectors are sorted by their contribution to reconstruction accuracy as shown in (c). **(c)** Cumulative reconstruction accuracy of the sorted basis vectors. **(d)** The basis vectors are classified as 1^st^, 2^nd^, or 3^rd^ order by convolving each with a simple image: a white circle on black background (left). Representative examples selected from the first 113 basis vectors are shown in the three boxes. *Left box*: basis vectors with 1^st^-order derivative characteristics in detecting the change in distance *D* (slope). *Middle box:* basis vectors with 2^nd^-order derivative characteristics in detecting the rate of change in distance *D* (curvature). *Right box:* basis vectors with 3^rd^-order derivative characteristics (change in curvature). **(e)** Cumulative count of basis vectors for each category shown in (d). **(f)** Reconstruction error of the basis vectors for each category identified in (d). The basis vectors in each category are sorted by their contribution to the reconstruction error.

To further assess the relevance of the different types of basis vectors, each category was separately tested for reconstruction accuracy. The 2^nd^-order category resulted in significantly smaller reconstruction error than either the 1^st^-order (Mann-Whitney rank test: U = 18042, p < 0.001) or 3^rd^-order categories (Mann-Whitney rank test: U = 1507754, p < 0.001). Unsurprisingly, reconstruction accuracy for the 1^st^-order category was by far the lowest of the three (Median = 0.003)

Within the first 113 leading basis vectors, the 1^st^-order category includes a total of 4 basis vectors, the 2^nd^-order category includes 62, and the 3^rd^-order category includes 47 basis vectors. Given that reconstruction accuracy typically correlates with the number of basis vectors, one possibility is that the differences in reconstruction error between categories may just be an artifact of the different number of basis vectors in each category. However, sorting the basis vectors in each category by their contribution to the reconstruction error, we find that there is an intrinsic difference in the amount of information carried by the basis vectors in each category. The results in Fig. 3f show that the reconstruction error drops the fastest with additional basis vectors for those in the 2^nd^-order category while the slowest for those in the 1^st^-order category. This result indicates that the 2^nd^-order basis vectors carry more information about the original patch than the 1^st^ or 3^rd^ order basis vectors. Thus, even though the number of basis vectors in each category does affect the reconstruction accuracy, the order of the basis vector plays a significant role in the observed reconstruction errors shown in Fig. 3f.

### 2.3. Tactile coding model based on curvature information

Sparse coding and ICA are powerful statistical models that seem to capture aspects of the coding strategies in visual as well as auditory cortices [4, 22, 28]. The sparse coding algorithm originally introduced by Olshausen and Field (1997) used data from natural images to estimate basis functions that had striking similarities to the receptive fields of simple cells in the primary visual cortex [45].

Because the ICA results identified curvature as a good candidate to represent 3D shape information, we used a similar approach and applied the FastICA algorithm to the 384-dimensional curvature maps ***P***(*C*) of natural objects. Examples of ***P***(*C*) used to obtaining the ICs are shown in Fig. 4a. The leading 100 basis vectors for the curvature map, sorted again by the method proposed in [44], are shown in Fig. 4b. In contrast to the results for the distance-based analysis (Fig. 3b), the basis vectors for the curvature map show localized Gabor-like properties similar to those found for natural images [22, 28]. This result is consistent with experimental work that identified oriented Gabor filters as good predictors for the filter properties of S1 neurons [46, 47].

**Fig. 4.**
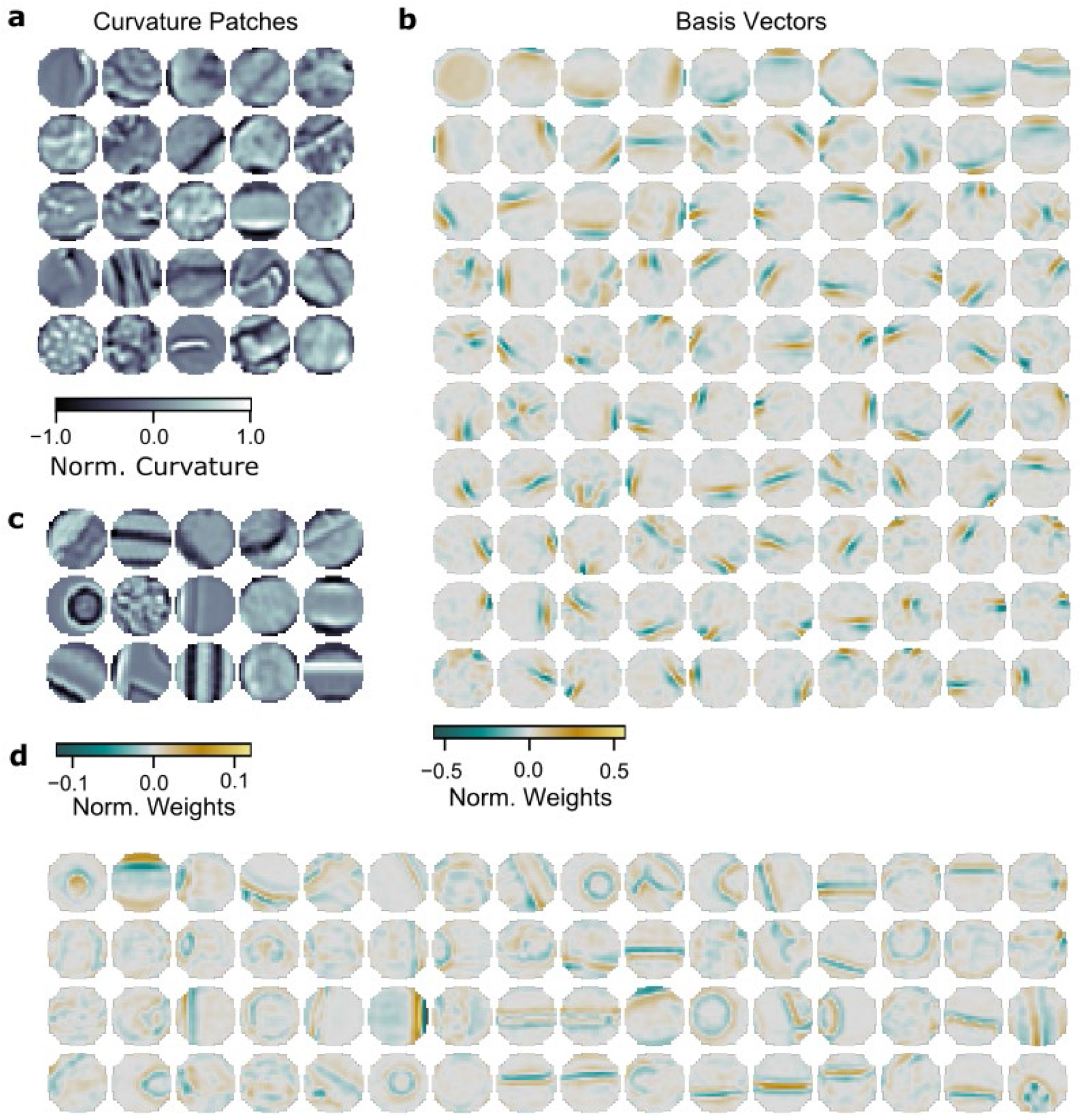
Basis vectors estimated by ICA for surface patches characterized by curvature *C.* **(a)** Example surface patches (*R* = 6 mm) randomly sampled from natural 3D data that show spatial maps of the curvature *C.* Grey scale is normalized to the maximum absolute value of the curvature within each patch. **(b)** The leading 100 basis vectors (columns of mixing matrix ***A***) as estimated by ICA from the curvature *C* of natural 3D data shown in (a). The mean of each basis vector was subtracted and the norm of the basis vectors was scaled to 1. **(c)** Same as (a) but for human-made 3D data. **(d)** Subset of 64 basis vectors (columns of mixing matrix ***A***) as estimated by ICA from curvature *C* of human-made 3D data shown in (c). The mean of each basis vector was subtracted and the norm of the basis vectors was scaled to 1.

Even more importantly, the same analysis applied to data for human-made objects (Fig. 4c) does not generate basis vectors with localized, Gabor-like patterns (Fig. 4d). Instead, basis vectors estimated from human-made objects have very regular characteristics, extending across the entirety of the patches and are even sometimes curved. Notably, these basis vectors seem to mimic the geometrical properties of the original curvature patches sampled from human-made objects (see Fig. 4c and 4d). Thus, although the first-order statistics of natural and human-made objects are similar (Fig. 2), these objects clearly differ in their second-order statistics.

### 2.4. Filter properties of S1 model neurons

To test whether the curvature-based basis vectors identified above might be associated with filter properties of S1 neurons, we used the neural model of Bensmaia et al. [46] to investigate the orientation selectivity of the corresponding ICA filters (rows of unmixing matrix ***W***). The proposed neural model [46] is described by the equation:

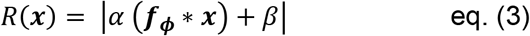

where ***x*** is the stimulus, *R*(***x***) is the neural response, ***f_ϕ_*** is a filter characterized by a set of parameters ***ϕ***, and *α* and *β* are scalar coefficients. The * operator symbolizes convolution; the parameters ***ϕ**, α*, and *β* are fit to model the response of each neuron. Bensmaia et al. [46] showed that Gabor filters for ***f_ϕ_*** accounted for 57% to 68% of the variance in individual neural responses.

To test our estimated filters, we generated model neurons by replacing ***f_ϕ_*** with each of the rows of ***W*** as estimated by ICA, and set *α* = 1 and *β* = 0. This procedure resulted in a population of 384 artificial neurons, each one based on one of the estimated ICA filters corresponding to the basis vectors for the curvature analysis of natural objects (the leading 100 of these filters are shown in Fig. 4b). To characterize the response of these artificial neurons we generated a set of bar stimuli with 16 different orientations between 0 and 180 degrees, similar to those used in the experiments of Bensmaia et al. (Fig. 5a). These stimuli, represented in terms of curvature, were used as input to the artificial neurons. For each artificial neuron, we measured the preferred direction, defined as the orientation resulting in maximum neural response, and the orientation selectivity, defined as the ratio of the neuron’s response in the preferred direction to its response in all other directions (see *Methods*).

**Fig. 5.**
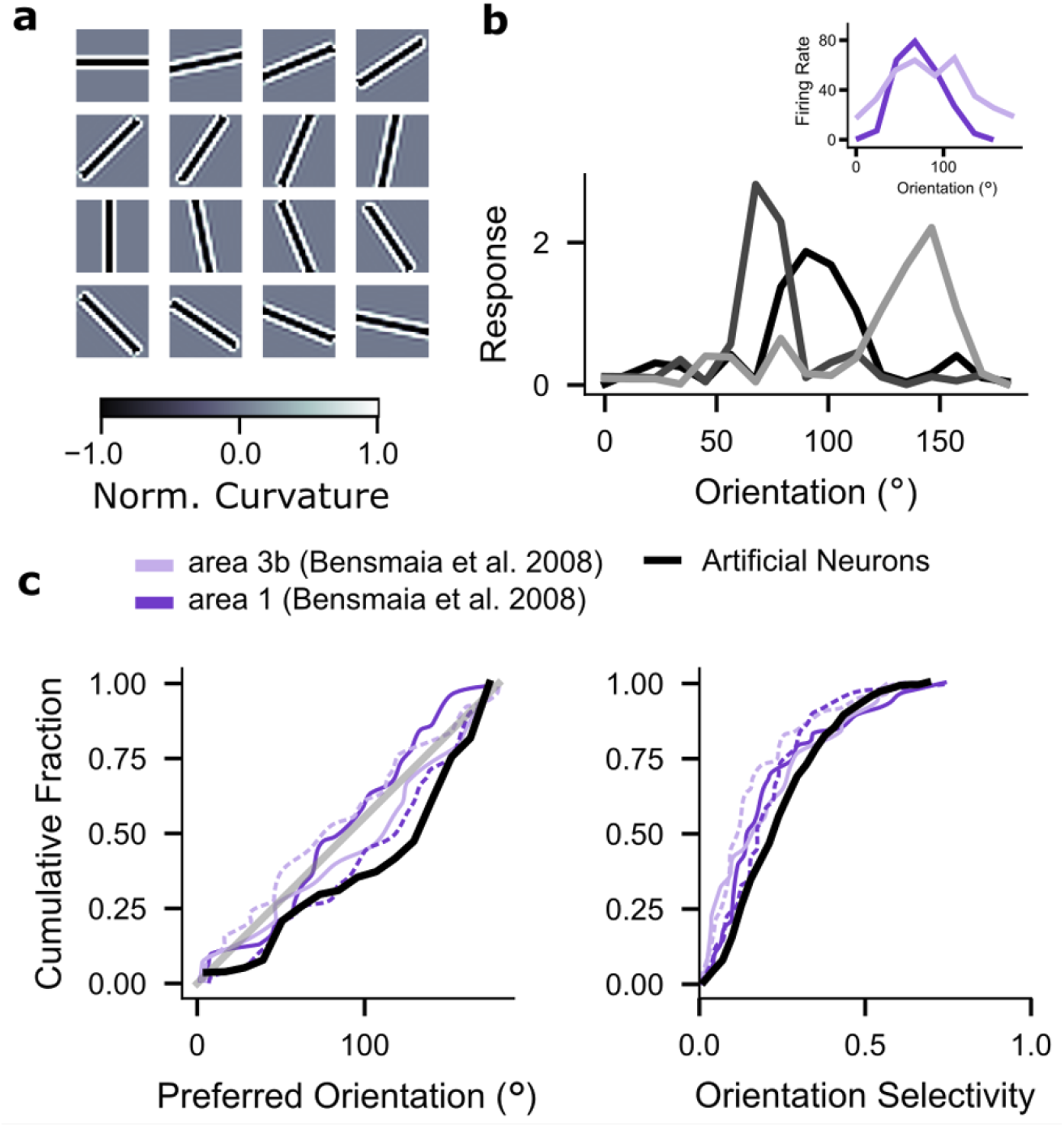
Orientation selectivity of model neurons based on ICA filters. **(a)** Curvature maps of the bar stimuli used to test artificial neurons based on ICA filters for the curvature analysis of natural objects. The orientation varies between 0 (top left) and 168.75 degrees (bottom right) in steps of 11.25 degrees. **(b)** Responses of three example artificial neurons with the highest orientation selectivity as a function of bar orientation in degrees. *Inset:* Example neurons in area 1 and 3b of S1 from experiments conducted by Bensmaia et al. [46]. **(c)** Cumulative distributions of neural responses across all 384 artificial neurons, each based on one of the ICA filters. *Left:* Cumulative fraction of artificial neurons as a function of their preferred orientation (solid black). Experimental results from area 1 and area 3b in S1 by Bensmaia et al. are shown in purple for bars indented into the fingertip (solid lines) and bars scanned over the fingertip (dashed lines). The 45-degree line representing a uniform distribution is shown in solid grey. *Right:* Same as *left* but cumulative fraction as a function of orientation selectivity.

Fig. 5b shows three artificial neurons that are strongly orientation selective (95^th^ percentile); they have preferred directions at approximately 65, 90, and 145 degrees. Examples of cortical neurons in area 1 and 3b from Bensmaia et al. [46] are shown in the inset for comparison. The cumulative fraction of the preferred orientation across all artificial neurons is shown in the left panel of Fig. 5c (black). The experimental results by Bensmaia et al. [46] are shown in purple, with solid lines indicating neural responses elicited by bars indented into the fingertip and dashed lines indicating neural responses activated by bars scanned across the fingertip. The selectivity of the artificial neurons closely approximates the experimental data in Bensmaia et al. [46]. In the right panel of Fig. 5c, we compare the cumulative fraction of orientation selectivity of the artificial neurons (black) with that of the recorded S1 neurons (purple, same as in left panel). Although the model neurons have not been fit to any neural data and result only from the statistical analysis of curvatures in natural objects, the cumulative fraction of orientation selectivity shows trends similar to those of neurons recorded in areas 1 and 3b in S1 [46].

## 3. Discussion

The statistical properties of visual scenes have been extensively studied, including studies of the global statistics of images in terms of luminance [6, 40, 48], contrast [6, 40, 49], and color [50, 51], as well as contours [52, 53], occlusion [52, 54–56], and eye movement [52]. However, all these studies primarily focus on the statistics of two-dimensional (2D) images, as proxies for 2D representations of the retinal image. Only a few attempts have been made to capture and describe the statistics of the 3D visual world [41].

For somatosensation, it is crucial to consider the 3D geometry of the environment with which the sensors interact. The work presented here has taken the first steps to quantify the statistics of 3D natural and human-made tactile scenes at the scale of human touch. The first-order statistics of local shape measurements (distance, slope, curvature) follow trends similar to those previously quantified for both camera and range images. Previous work showed that both these types of images exhibit different first-order statistics for natural (vegetation) and human-made (urban) scenes, mainly showing higher kurtosis for human-made scenes [55]. Other work showed that spatial low-frequency components are much more prominent in human-made scenes than in natural scenes [57]. We observed similar trends in our data.

The ICA analysis of higher order statistics reveals that basis vectors with second-order filter characteristics contribute significantly to accurate reconstruction of the data. These results suggest that curvature is an important metric to represent the 3D shape of objects at the spatial scale of a fingertip. Previous experimental work supports this hypothesis. Experiments in monkeys have shown that the spike rate of cutaneous mechanoreceptors in the finger pad respond to the amount of curvature and its rate of change, both for slowly adapting and rapidly adapting afferent fibers [33, 36, 38]. In addition, areas in human somatosensory cortex have been found to be active during a shape discrimination task that specifically tested for surface curvature [58]. A study in macaque inferior temporal cortex found neurons that are tuned to the amount of curvature and to curvature direction [59]. Correlations between curvature direction and neural activity were also found in a study that examined shape coding in visual V4 and somatosensory cortices, suggesting that these areas use similar mechanisms to encode shape [60].

The ICA of natural curvature data revealed basis vectors with Gabor-like characteristics similar to those found for natural images [45]. Further analysis with a simple model based on artificially generated neurons that use the IC filters showed neural responses that accurately capture the statistics of S1 neural responses in primates [46]. These results suggest that efficient coding algorithms such as ICA may yield insights about the filter properties of cortical neurons that go beyond those identified by receptive field measurements [46].

Although the present analysis tactile scenes has focused on the scale relevant to human touch, the methods used are not restricted to that scale or to a specific strategy for sampling the environment. Our approach can be extended to patches of different sizes explored with different degrees of spatial resolution. As such, the data presented in this study is best considered as a small subset of the data that an organism might acquire over a lifetime or even over generations. Another important extension to the framework developed here is to incorporate the temporal component that is crucial to tactile systems. Movement allows animals to gather information over time, as tactile sensors are moved over a surface. Here we have limited our analysis to a static component that describes the aggregated tactile information obtained over such a sweep by concentrating on the intrinsic properties of object patches.

This work examined the statistics of three metrics: distance, slope, and curvature, as intrinsic properties of external objects; patches were randomly sampled from each objects’ surface. However, animals and humans sample the world with a goal in mind (such as identifying an object) and combine past and current sensory information both to predict future sensory input as well as increase search efficiency. Therefore, we expect that subjects would sample a set of patches that differs from the randomly sampled ones used here. This hypothesis could be tested by asking human subjects to identify objects based only on tactile input in a situation where the object is behind a screen and thus visually inaccessible. To approximate the conditions of the work presented here, the subject would be allowed to use only one finger to touch the surface of the objects. An efficient exploration could be enforced by limiting the time available for object identification. The objects would be the same ones used in the presented study; here we have characterized the statistics P(W) of these objects as representative of the world W. In the experiment with human subjects, the places on the object’s surface explored by the subject as they try to identify the object would be tracked. This data would allow for a characterization of the statistics of the world when sampled by volitional exploration E. This P(W|E) could then be compared to the P(W) analyzed here.

## 4. Methods

### 4.1. Data acquisition

We used a *EinScan Pro 2X* scanner to capture 3D point cloud representations of the surface of both natural objects and human-made objects. The scanner was mounted on a *Shining 3D* desktop tripod while the objects were placed on the *Shining 3D* turntable. The *EXScan Pro* software was used to capture the scans in “Fixed Scan” mode as “Non-Texture Scans” (without color information). For each scan, the turntable performed 30-50 steps for one rotation and one rotation per scan. Typically, each object was scanned in 2-3 different orientations and the scans were then combined into a single point cloud. One of the objects was very small in size and was therefore scanned using “Handheld Rapid Scan” mode with highest resolution.

### 4.2. Data cleaning

Each point cloud was imported to *Geomagic Design X* software for cleaning and meshing. First, the standard meshing algorithm of *Geomagic Design X (Mesh Buildup Wizard™)* was used to triangulate the point cloud. Outlier data (non-manifold triangles, small clusters, and isolated triangles) were removed manually or using the *Healing Wizard* software. Holes in the scan were filled using the *Fill Holes* tool. For larger holes or for holes at the boundary, the *Add Bridge* tool was used to segment the holes to ensure better reconstruction with the subsequent use of the *Fill Holes* tool. After all holes were filled and unusable parts of the mesh had been removed, the scan was remeshed targeting an average edge length of 0.1mm. After remeshing, the mesh was enhanced (smoothed and sharpened) with medium settings and optimized to improve curvature flow at medium settings with a maximum of 10 iterations. Finally, the centroid of the mesh was redefined to be the origin and exported in *Binary STL* format, which stores each triangle as a collection of three edge points and the corresponding face normal. The resulting dataset consisted of 137 scans, of which 96 represented natural scenes and 41 represented human made objects (SI Dataset).

### 4.3. Sampling and computation of shape metrics

The method used to compute the distance (*D*), slope (*S*), and curvature (*C*) metrics is illustrated in Fig. 6. First, we randomly sampled 100-150 circular surface patches of radius *R* from each object (Fig. 6a). The point cloud for each sampled surfaced patch was analyzed using Principal Component Analysis (PCA). The plane spanned by the two leading PCs was found, and the points in the surface cloud were projected onto this reference plane (Fig. 6b). The location of the projected points within the plane was digitized to a 24×24 grid (Fig. 6c). The position of the grid cell in the *i*^th^ row and *j*^h^ column is denoted as *p_i,j_*; the width of the grid cells is *δ* = *R*/12.

**Fig. 6.**
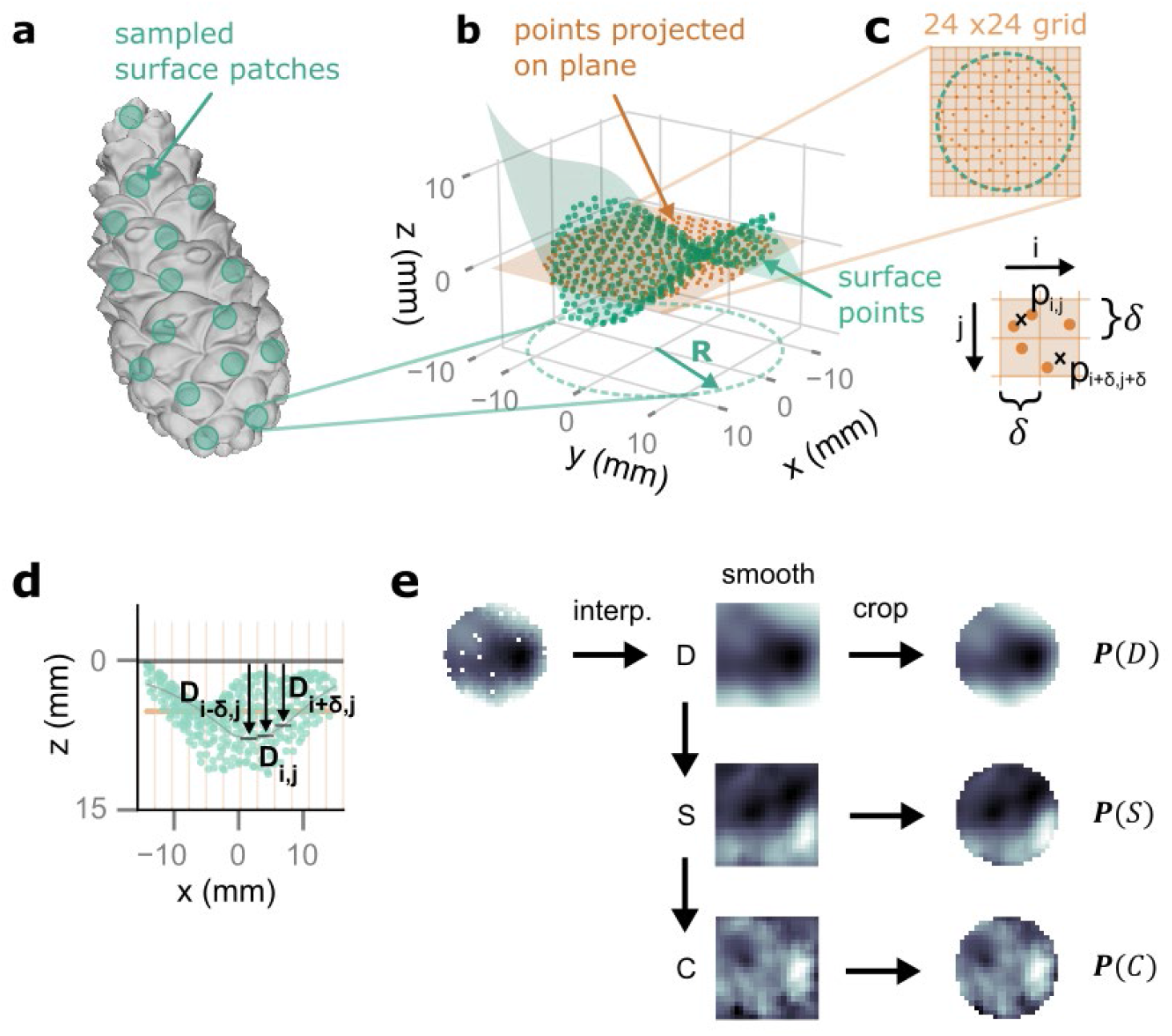
Processing of 3D data. **(a)** Approximately 100-150 circular surface patches of radius *R* were randomly sampled from each object scan. **(b)** For each surface patch, the average plane of the point cloud (orange) was found by using Principal Component Analysis (PCA). **(c)** The points in the cloud were then projected onto the plane, and their location within the plane digitized to a 24×24 grid. The position of the grid cell in the *j*^tj^ row and *i*^th^ column was indicated as *p_i,j_*. The cells are of linear size *δ* = *R*/12. **(d)** For each cell *(i,j)* the distance *D_i,j_* to the reference plane describes the local shape of the patch. **(e)** The distances *D* were interpolated to the entire grid, smoothed, and cropped to a circular patch. The metrics slope *S* and curvature *C* were computed from the interpolated and smoothed grid before cropping it to a circular patch.

The plane was then translated in the direction of its outward normal vector, away from the surface of the object just until all surface sampled points were below it. For each point (*i, j*) on the surface cloud, the distance *D_i,j_* to the reference plane was computed as the orthogonal distance from that point to its projection (Fig. 6d). The distances *D* were then interpolated over the entire grid, smoothed, and cropped to a circular patch of radius *R* to obtain a map ***P***(*D*) over 448 grid points. Patches that filled less than 75% of the circular area before interpolation were rejected.

The metrics *S* and *C* were computed from the interpolated and smoothed square grid before cropping it to a circular patch. The slope S was calculated as the forward difference in *D* between neighboring points in the grid (Eq. 1), resulting in a circular map of 421 slope values ***P***(*S*). Similarly, *C* was calculated as the second-order central difference in *D* (Eq. 2), resulting in a circular map of 384 curvature values ***P***(*C*). The resulting maps for one example patch are shown in Fig. 6e.

Since the square grid size is fixed at 24×24 before cropping it to a circular patch, the spatial resolution of the resulting circular maps is determined by the patch radius *R*. Each circular patch can be viewed as a sensory surface with a fixed number of receptors represented by the grid. A larger *R* leads to a grid that covers a larger area sampled at a lower spatial resolution. Based on the average contact area of a finger pad (~120 mm^2^ at 1N normal load [39]), we chose *R* to be 6 mm (corresponding to 113 mm^2^ of circular contact area) for the results presented here. The choice of a grid size of 24×24 is based on the average edge length of the 3D meshes of the data (~0.1 mm) rather than on biologically plausible receptor density. With an *R* of 6 mm, the datasets for natural and human-made objects consisted of a total of 12,309 and 5,788 patches, respectively.

Because the slope of a surface depends on the direction along which it is taken, its value for a given grid cell was computed by averaging the slope in the x direction 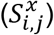 and the slope in the y direction 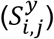. The equations for 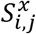 and 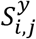 are:

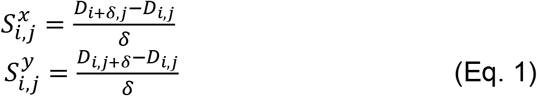

The curvature of a surface is also direction dependent, and was again computed as the average of the curvature in the x direction 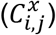 and the curvature in the y direction 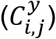. The equations for 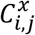 and 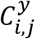 are:

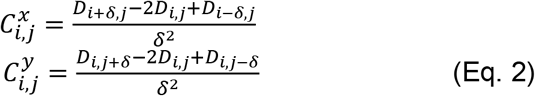

### 4.4. Neural model

The orientation selectivity *os* is computed as:

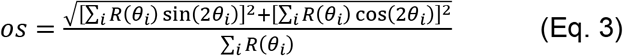

where *R*(*θ_i_*) represents the response of the artificial neuron to a bar stimulus with orientation *θ_i_*. Stimulus orientation ranged from 0 to 180 degrees in intervals of 11.25 degrees, for *i* = {1,2, …16}. The orientation selectivity of a neuron ranges from 0 to 1, where 0 indicates a uniform response to all orientations and 1 indicates that the neuron responds only to a single stimulus orientation. For more details about this measure of orientation selectivity see [46].

